# DisCovER: distance- and orientation-based covariational threading for weakly homologous proteins

**DOI:** 10.1101/2020.01.31.923409

**Authors:** Sutanu Bhattacharya, Rahmatullah Roche, Debswapna Bhattacharya

## Abstract

**Motivation:** Threading a query protein sequence onto a library of weakly homologous structural templates remains challenging, even when sequence-based predicted contact or distance information is used. Contact- or distance-assisted threading methods utilize only the spatial proximity of the interacting residue pairs for template selection and alignment, ignoring their orientation. Moreover, existing threading methods fail to consider the neighborhood effect induced by the query-template alignment.

**Results:** We present a new distance- and orientation-based covariational threading method called DisCovER by effectively integrating information from inter-residue distance and orientation along with the topological network neighborhood of a query-template alignment. Our method first selects a subset of templates using standard profile-based threading coupled with topological network similarity terms to account for the neighborhood effect and subsequently performs distance- and orientation-based query-template alignment using an iterative double dynamic programming framework. Multiple large-scale benchmarking results on query proteins classified as hard targets from the Continuous Automated Model Evaluation (CAMEO) experiment and from the current literature show that our method outperforms several existing state-of-the-art threading approaches; and that the integration of the neighborhood effect with the inter-residue distance and orientation information synergistically contributes to the improved performance of DisCovER.

**Availability:** https://github.com/Bhattacharya-Lab/DisCovER

## 1 Introduction

Accurate prediction of the three-dimensional (3D) structure of a protein from its sequence remains challenging (Dill and MacCallum, 2012). Template-based modeling (TBM), one of the most reliable and accurate approaches for protein 3D structure prediction, uses homologous templates of analogous folds available in the Protein Data Bank (Berman *et al*., 2000) to infer 3D structural models of unknown proteins. As such, the success of TBM intrinsically depends on the detection of homologous templates and the generation of accurate query-template alignments. However, in the absence of close sequence homology, template detection and alignment become challenging. Protein threading is a TBM approach that can address the challenge by leveraging multiple sources of information such as sequence profile, secondary structure, solvent accessibility, and torsion angles. Traditional threading methods such as HHpred (Söding, 2005), MUSTER (Wu and Zhang, 2008), SparkX (Yang *et al*., 2011), pGenThreader (Lobley *et al*., 2009), and CNFpred (Ma *et al*., 2012, 2013) have shown noteworthy success by successfully modeling protein structures even in the absence of significant sequence homology. Nevertheless, the performance of traditional threading methods sharply declines when the evolutionary relationship between the query and templates becomes very low (Zheng *et al*., 2019), the so-called remote-homology threading scenario.

With significant recent progress in residue-residue contact or distance prediction mediated by sequence co-evolution and deep learning (Jones *et al*., 2012; Kamisetty *et al*., 2013; Jones *et al*., 2015; Li *et al*., 2019; Greener *et al*., 2019; Wang *et al*., 2017; Xu, 2019; Yang *et al*., 2020), predicted contact or distance information becomes a valuable source of additional information for remote-homology threading. Consequently, predicted contact- or distance-based threading has attracted promising attention. For instance, EigenTHREADER (Buchan and Jones, 2017) utilizes contacts predicted by MetaPSICOV (Jones *et al*., 2015) to perform contact map threading. map_align (Ovchinnikov *et al*., 2017) employs iterative double dynamic programming driven by co-evolutionary contact predictor, GREMLIN (Kamisetty *et al*., 2013). Very recently, CEthreader (Zheng *et al*., 2019) and CATHER (Du *et al*., 2020) perform contact-assisted threading using contacts predicted by ResPre (Li *et al*., 2019) and MapPred (Wu *et al*., 2020), respectively, together with sequence profile-based features. DeepThreader (Zhu *et al*., 2018) goes one step further by incorporating finer-grained distance information instead of contacts for boosting threading performance (Xu and Wang, 2019).

While these methods exploit predicted contact or distance information during threading often in conjunction with sequential information, they do not consider two key factors that can further improve threading accuracy. First, the recent extension of deep residual network architecture has resulted in accurate inter-residue orientations predicted from coevolution (Yang *et al*., 2020) in addition to distances, but none of the threading methods incorporate orientation information. Second, most of the threading approaches do not include the effect of the residue pairs in the neighborhood of an aligned query-template residue pair. That is, they ignore the neighborhood effect induced by the query-template alignment.

In this article, we introduce a new distance- and orientation-based threading method DisCovER (<u>Dis</u>tance- and orientation-based <u>Cov</u>ariational thread<u>ER</u>) that effectively integrates information from inter-residue distance and orientation along with the topological network neighborhood of a query-template alignment using an iterative double dynamic programming framework to greatly improve threading template selection and alignment. Experimental results show that our new method performs better than profile-based threading methods SparkX, HHpred, CNFpred, MUSTER, PPAS, and pGenThreader; as well as state-of-the-art contact-assisted approaches CEthreader, map_align, Eigen-THREADER, and CATHER on weakly homologous proteins. At one of the most challenging threading situations, DisCovER yields better performance than the RaptorX server (Källberg *et al*., 2012; Zhu *et al*., 2018) participating in the Continuous Automated Model Evaluation (CAMEO) experiment (Haas *et al*., 2019) and employing the distance-based threading method DeepThreader (Zhu *et al*., 2018). DisCovER is freely available at https://github.com/Bhattacharya-Lab/DisCovER.

## 2 Materials and methods

### 2.1 Feature sets and inter-residue geometries

Our feature set includes both sequential and pairwise features for the query protein and templates. For a query protein, we generate sequence profiles based on multiple sequence alignments (MSA) (Zhang *et al*., 2020), and predict profile-based features including secondary structure, solvent accessibility, and backbone dihedral angles using SPIDER3 (Heffernan *et al*., 2017). We also predict inter-residue geometries including distances and orientations by feeding the MSAs into trRosetta (Yang *et al*., 2020). The predicted distance map is then binned into 9 segments at 1Å interval: <6Å, <7Å, …, <14Å, by summing up probabilities for distance bins below specific distance thresholds. The predicted inter-residue orientations include two dihedral angles (ω, θ), both binned into 24 evenly spaced segments with a bin width of 15° each, and one planar angle (φ), binned into 12 evenly spaced segments with a bin width of 15° each. All distance orientation bins have associated likelihood values for the query protein predicted by trRosetta. For the templates, we use structure-derived profiles, extract secondary structures and solvent accessibility using DSSP (Kabsch and Sander, 1983), and compute backbone dihedral angles, inter-residue distance maps, and orientation information including ω, θ dihedrals and φ angle from the structure.

### 2.2 Geometry-based scoring of a query-template alignment

DisCovER scores a query-template alignment as follows:

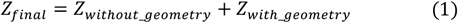

where *Z*_*final*_ is the normalized alignment score for selecting the top-ranked template, *Z*_*without_geometry*_ is the normalized alignment score based only on profile information with neighborhood effect, and *Z*_*with_geometry*_ is the normalized alignment score using profile information, neighborhood, and inter-residue geometries including distance and orientation. In the following, we describe each term in detail.

### Stage 1 Scoring profile-based alignment with neighborhood effect

A profile-based query-template alignment is scored for aligning the *i*th residue of the query and the *j*th residue of the template similar to our recent work (Bhattacharya and Bhattacharya, 2019) as follows:

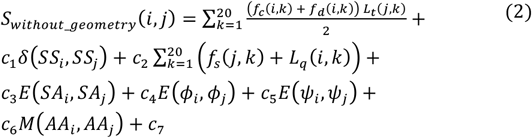

where the first term defines the sequence profile-profile alignment. *f*_*c*_(*i, k*) and *f*_*d*_(*i, k*) define the frequency of the *k*th residue at the *i*th query position of the MSA for “close” and “distant” homologous sequences, respectively. The frequency is determined using the Henikoff weighting scheme (Henikoff and Henikoff, 1994). *L*_*t*_ is the log-odds profile of the template for the *k*th residue at the *j*th position, which is obtained by PSI-BLAST (Altschul *et al*., 1997) with an E-value of 0.001. The next term measures the consistency between the predicted and observed three-state secondary structures, such that the function δ returns 1 if two variables are matched and −1 otherwise. The next term is the agreement between the structure-derived profiles (*f*_*s*_) of the *k*th residue at the *j*th position of the template structure and the sequence profile (*L*_*q*_) of the *k*th residue at the *i*th position of the query sequence. The function *E* in the next three terms is defined by: *E*(*x*_*i*_, *x*_*j*_) = (1 − 2|*x*_*i*_ *– x*_*j*_|), where the variables are the predicted and the observed values of relative solvent accessibility (*SA*) and backbone dihedral angles (*ϕ, ψ*) of the *i*th position of the query and the *j*th position of the template, respectively. The seventh term corresponds to the match between the hydrophobic residues of the query and the template. c is the weighting parameter adopted from (Bhattacharya and Bhattacharya, 2019).

To further improve the sensitivity of profile-based alignment, we borrow ideas from comparative network analysis. We adopt an approach similar to that was originally used in the IsoRank network alignment algorithm (Singh *et al*., 2008) and very recently adopted in network-based structural alignment of RNA sequences (Chen *et al*., 2019), in which two nodes in different networks are more likely to be aligned to each other if their neighbors are also aligned well to one another. This results in a similarity diffusion scheme to compute the agreement between the networks, leading to an improved alignment. Following similar principles, we integrate *connected similarity*, attempting to estimate the topological agreement between the query and the template by capturing the similarity between the neighborhoods of two residues.

### Connected similarity

(*S*_*c*_) is based on the principle that one query-template residue pair is likely to be aligned if their neighboring residues are also aligned. It is calculated for the residue pair (*i, j*) as follows:

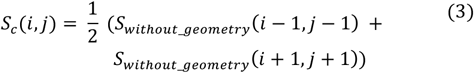

Connected similarity (*S*_*c*_) is then added to the profile-based alignment score as follows:

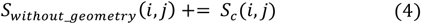

We use the Needleman-Wunsch (Needleman and Wunsch, 1970) global dynamic programming to score every query-template alignment. To select the top-fit templates, we compute the *Z*_*without*_*geometry*_ based on the raw alignment score *S*_*without*_*geometry*_ to assess the quality of each query-template alignment as follows:

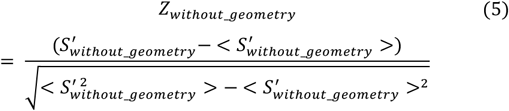

where 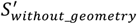 is the larger one of the raw alignment score *S*_*without*_*geometry*_ divided by the full alignment length (including query and template ending gaps) and the partial alignment length (excluding query ending gaps). 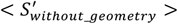 is the average 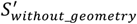 across all templates in the template library. A subset of fifty top-scoring templates is selected for the next stage.

### Stage 2 Scoring distance- and orientation-based alignment

A similarity score is calculated for each row of query (Q) and template (T) distance maps as follows:

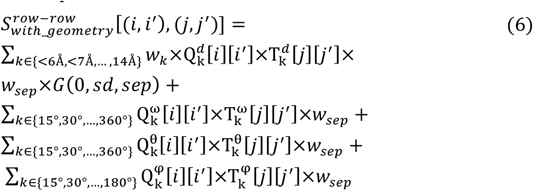

where the first term calculates the similarity between the predicted distance map of the query and true distance map of the template at a given distance threshold of *k*, where *k* ∈{<6Å, <7Å, …, <14Å}. 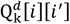 is the predicted likelihood value of the residue pair *i* and *i*′ of the query to be within a distance threshold of *k*Å;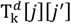 is a Heaviside step function that has a value 1 if the residue pair *j* and *j*′ of the template is within the distance threshold of *k*Å and 0 otherwise; w_*k*_ is the corresponding weight parameter adapted from the literature (Du *et al*., 2020) with 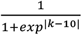; *w*_*sep*_ is the weight of the minimum of sequence separation of query residues and template residues defined as *w*_*sep*_=0.75 for separation= 5 and log10(1+separation) for separation ≥ 6, similar to other studies (Ovchinnikov *et al*., 2017; Greener *et al*., 2019); *G*(0, *sd, sep*) is a zero-mean Gaussian function, which is also adopted from the literature (Ovchinnikov *et al*., 2017) and defined as *exp*(-*sep*^2^/(2 *sd*^2^)), where *sep* is an absolute difference of sequence separation of query residues and sequence separation of template residues, and *sd* or standard deviation is a function of the smaller of the sequence separation of query residues and the sequence separation of template residues . The next three terms calculate dihedral (ω, θ) and planar (φ) angles similarities between the query and template residue pair. We treat the orientation information similar to distances and compute the similarity between the predicted ω, θ dihedrals or φ angle of the query and the corresponding true angles of the template at a specific angle bin of *k*, where *k* ∈{15°,30°,…,360°} for the ω, θ dihedrals and *k* ∈{15°,30°,…,180°} for the φ angle. Analogous to distances, 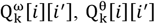, and 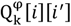 are the predicted likelihood values of the residue pair *i* and *i*^°^ of the query to be within an angle bin of *k*° for the angles ω, θ, and φ, respectively. Similarly, 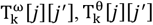, and 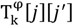 are the Heaviside step functions corresponding to the angles ω, θ, and φ, respectively, having values 1 if a residue pair *j* and *j*′ of the template is within an angle bin of *k*° and 0 otherwise. *w*_*sep*_ is the weight term described before.

Our double dynamic programming framework for computing the optimal alignment score between the query and each of the fifty top-scoring templates selected from the previous stage comprises of two dynamic programming steps. In the first step, we perform row-by-row comparisons between the query and template. Dynamic programming is used to find the alignment for the two rows being matched which maximizes the composite distance- and orientation-based alignment score described in equation 6. These scores are stored in a similarity matrix and are used to obtain the optimal alignment by using the Smith-Waterman (Smith and Waterman, 1981) algorithm. At this step, however, the scores for individual row-row comparisons are overestimated since the alignments for each pair are independently computed in the first step. We subsequently update the similarity matrix using a second step based on the current alignment by employing a second dynamic programming. Such an iterative updating strategy is originally proposed in (Taylor, 1999) and later adopted in (Ovchinnikov *et al*., 2017), although our score is quite different. After obtaining the optimal alignment from the similarity matrix, the profile score and the gap-score are re-calculated to compute the raw alignment score (*S*_*with_geometry*_). The similarity score for each query-template pair is normalized using *Z*_*with_geometry*_ as follows:

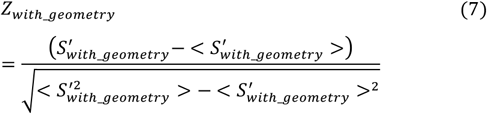

where 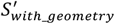 is the larger one of the raw alignment score *S*_*with_geometry*_ divided by the full alignment length (including query and template ending gaps) and the partial alignment length (excluding query ending gaps). ⟨…⟩ denotes the average 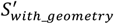 of all top-scoring templates.

### Building full-length 3D models

After selecting the first-ranked template using Equation (1), we use MODELLER (V9.22) (Webb and Sali, 2014) to generate the full-length 3D model of a query protein using the associated query-template alignment. In addition to employing the standard automodel() class of MODELLER for model building purely through the satisfaction of spatial restraints from the query-template alignment, we also experiment with model building with additional restraints from predicted distances, orientations, and secondary structures by redefining the automodel.special_restraints() class. Specifically, we feed bounded harmonic restraints for the predicted distance thresholds corresponding to the 9 distance bins used in query-template alignment with a minimum likelihood cutoff of 0.85 using the physical.xy_distance() function, bounded harmonic restraints for the predicted orientation information derived from the highest likelihood bins having the minimum likelihood cutoff of 0.85 with the ψ angle using physical.angle() function and ω, θ dihedrals using physical.dihedral() function, and secondary structure restraints for realizing the predicted secondary structure using the secondary_structure() module. Of note, all of these additional restraints are integrated to the list of spatial restraints derived from the query-template alignment to instruct MODELLER to satisfy them as best as it can.

### 2.3 Benchmark datasets, methods to compare, template libraries used, and threading performance evaluation

To evaluate remote-homology threading performance, we benchmark our new method DisCovER using targets from the Continuous Automated Model Evaluation (CAMEO) experiments consisting of 117 proteins classified as “hard” (Haas *et al*., 2019), released between 8 December 2018 and 1 June 2019 having length between 50 and 500 residues (range is 51 to 487). On this dataset, DisCovER is compared against profile-based threading methods such as SparkX (Yang *et al*., 2011), CNFpred (Ma *et al*., 2012, 2013), MUSTER (Wu and Zhang, 2008), PPAS (Wu and Zhang, 2007), and pGenThreader (Lobley *et al*., 2009); as well as state-of-the-art contact-assisted methods including CEthreader (Zheng *et al*., 2019) utilizing ResPRE-predicted contact maps (Li *et al*., 2019) and EigenTHREADER (Buchan and Jones, 2017) utilizing DMPfold-predicted maps (Greener *et al*., 2019). The template libraries for DisCovER, CEthreader, EigenTHREADER, SparkX, MUSTER, PPAS, pGenThreader are generated from the same set of 70,670 templates downloaded from https://zhanglab.ccmb.med.umich.edu/library/ (Yang *et al*., 2015), curated before the release of the CAMEO targets. The template library for CNFpred is downloaded from http://raptorx.uchicago.edu/download/. For all methods, we evaluate threading performance by comparing the top-ranked full-length predicted 3D models, built using the standard automodel() class of MODELLER from the query-template alignment, against the experimental structures of the target proteins using the TM-score metric (Zhang and Skolnick, 2004), which ranges from 0 to 1 with higher score indicating better performance and TM-score >0.5 indicating the attainment of correct overall fold (Xu and Zhang, 2010). Of note, DisCovER utilizes distances and orientations predicted from trRosetta (Yang *et al*., 2020), which uses a training set collected from a snapshot as of 1 May 2018, older than the CAMEO test set used here. DisCovER also relies on secondary structure predictor SPIDER3 (Heffernan *et al*., 2017), which uses a much older training dataset and thus independent of the CAMEO test set. We collect publicly available multiple sequence alignments (MSAs) independently generated using non-overlapping protein sequence databases from https://yanglab.nankai.edu.cn/trRosetta/benchmark/ to feed into trRosetta and SPIDER3. Furthermore, the template library used in DisCovER excludes any templates released after starting of CAMEO experiments (8 December 2018), free from any overlap. Finally, the “hard” target difficulty classification of the CAMEO test set defined by CAMEO (Haas *et al*., 2019) warrants their non-overlapping and weakly homologous nature, thereby enabling us to focus on difficult targets in which existing methods have limitations (Zhu *et al*., 2018).

The definition of “hard” can be made even more stringent by requiring that TM-score of the HHpredB server (Söding, 2005) participating in CAMEO to be less than 0.5. This reduces the number of targets to 60. This harder set simulates one of the most challenging threading situations while enabling a comparison between DisCovER and the distance-based threading method DeepThreader (Zhu *et al*., 2018). DeepThreader method is not publicly available, but the RaptorX server (Källberg *et al*., 2012; Zhu *et al*., 2018) participates in CAMEO and employs the DeepThreader method according to the CAMEO assessment paper (Haas *et al*., 2019). While RaptorX uses PDB90 as the template database and builds 3D models using Rosetta (Xu and Wang, 2019) as opposed to MODELLER, we compare the performance of DisCovER on this common set of 60 very hard targets to that of DeepThreader after downloading the predictions submitted by the RaptorX server from the official website of CAMEO (https://cameo3d.org/) and computing their TM-scores.

We are unable to directly compare DisCovER with two other state-of-the-art contact-assisted methods map_align (Ovchinnikov *et al*., 2017) and CATHER (Du *et al*., 2020) on the CAMEO test set, because map_align is too computationally expensive to run locally given our limited computational resources and CATHER is only available as a webserver and thus not suitable for large-scale benchmarking. However, the published work of CATHER reports the mean TM-scores of 3D models predicted using various threading methods including CATHER, map_align, EigenTHREADER, HHpred (Söding, 2005), SparkX, and MUSTER over a dataset of 131 hard targets with pairwise sequence identity <25% and length ranging from 50 to 458 residues (Du *et al*., 2020). We use this set to compare DisCovER against CATHER and map_align by running DisCovER locally after excluding templates with sequence identity >30% to the query proteins, and comparing its average performance against the reported results of CATHER and map_align, in addition to the other threading methods presented.

## 3 Results

### 3.1 Performance on 117 hard targets from CAMEO

Over the 117 hard targets from CAMEO, our distance- and orientation-based threading method DisCovER performs better than the two contact-assisted threading methods as well as all five profile-based approaches. As shown in **Table 1**, DisCovER attains a mean TM-score of 0.505, which is higher than the next best contact-assisted threading method CEthreader (0.483) and the best among profile-based threading methods CNFpred (0.464). The performance improvement for DisCovER is statistically significant at 95% confidence level compared to all other methods. DisCovER also predicts higher number of correct folds (TM-score >0.5) with a success rate of 57.3%, which is ∼5% higher than the success rate of CEthreader and ∼9% higher than that of CNFpred. Of note, the best contact-assisted threading method CEthreader falls short of achieving a mean TM-score of 0.5, whereas DisCovER exceeds this criterion.

**Table 1.** Benchmark results of various threading methods on 117 hard targets from CAMEO.

| Methods | TM-score | *p*-value | #Correct folds |
| --- | --- | --- | --- |
| pGenThreader | 0.423 | 1.1E-08 | 46 |
| PPAS | 0.456 | 9.3E-05 | 54 |
| MUSTER | 0.459 | 0.0006 | 54 |
| SparkX | 0.461 | 0.0003 | 57 |
| CNFpred | 0.464 | 0.004 | 56 |
| EigenTHREADER | 0.461 | 1.9E-05 | 58 |
| CEthreader | 0.483 | 0.029 | 61 |
| DisCovER | **0.505** | - | **67** |

Column “*p*-value” represents one sample *t*-test’s *p*-value of the TM-score difference compared to DisCovER. Column “#Correct folds” represents the number of models with TM-score >0.5. Values in bold represent the best performance.

**Figure 1** shows the head-to-head comparison between DisCovER and the best contact-assisted threading method CEthreader and the best profile-based threading method, CNFpred. DisCovER attains higher TM-score for 74 and 66 targets compared to CEthreader and CNFpred, respectively. In summary, the advantage of DisCovER in threading remote-homology proteins over the others is significant.

**Fig 1.**
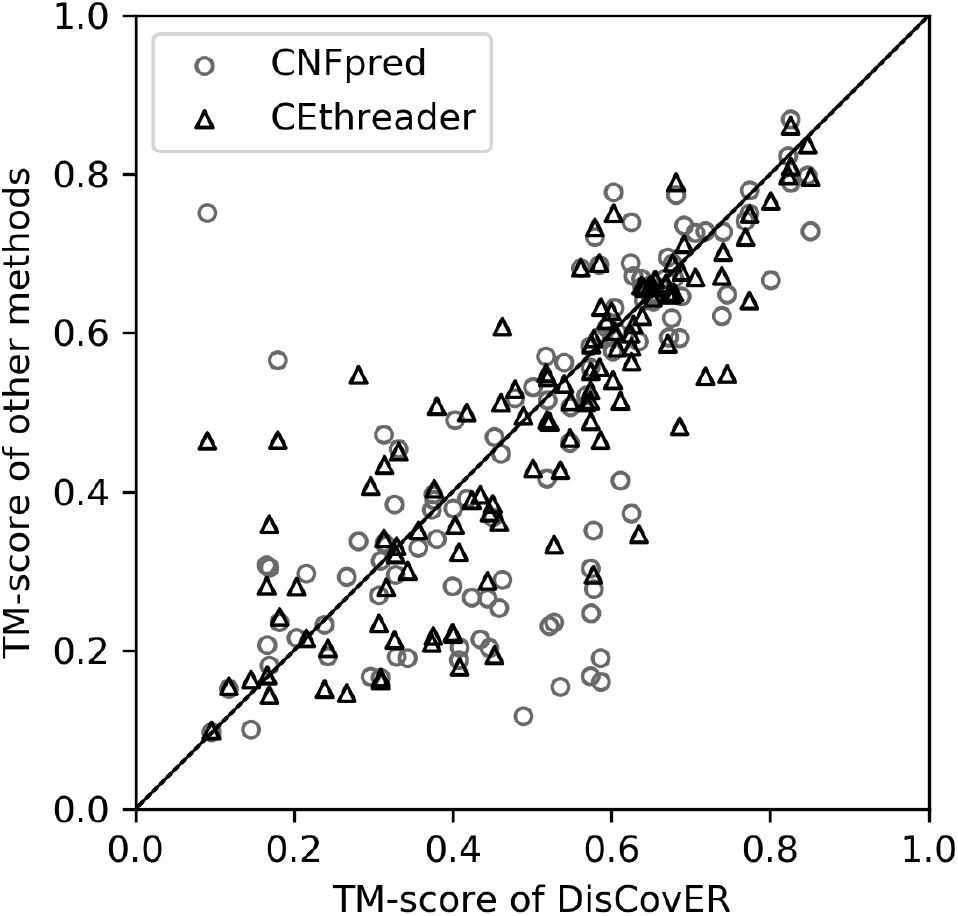
Head-to-head performance comparison between DisCovER (x-axis) vs. CEthreader and CNFpred (y-axis) on 117 hard targets from CAMEO.

### 3.2 Contribution of individual components

In addition to profile-based alignment, DisCovER incorporates a) neighborhood effect, b) distance, and c) orientation. To investigate the contributions of each of these components to DisCovER performance, we perform an ablation study on the CAMEO test set by gradually removing each component from the full-fledged DisCovER method one at a time and evaluate the performance. The same template library, sequence databases, and same set of features are used in all cases to generate the query-template alignments that are then fed into the standard automodel() class of MODELLER to generate the 3D structures.

As reported in **Table 2**, the full-fledged DisCovER attains the best performance (mean TM-score of 0.505) than any of its ablated variants over the 117 hard targets from CAMEO. Without orientation, the mean TM-score decreases to 0.503, which is further decreased to 0.488 without distance information. The performance of the ablated variant of DisCovER that only incorporates distance but no orientation outperforms the variant that only incorporates orientation but no distance. It is interesting to note that the average performance of either variant is still better than state-of-the-art contact-assisted methods CEthreader and EigenTHREADER, indicating that the incorporation of either distance or orientation information in DisCovER is sufficient to outperform top contact-assisted threading method such as CEthreader. When both distance and orientation terms are excluded, the mean TM-score drops to 0.472, but it is still better than the top profile-based threading method CNFpred. Upon further exclusion of neighborhood effect, the mean TM-score slightly reduces to 0.470. The results demonstrate that all components contribute to the improved performance of DisCovER, with distance and/or orientation information having significant contribution. We note that DisCovER incorporates distance and orientation information only in stage 2 for computing the optimal alignment score between the query and each of the fifty top-scoring templates selected from stage 1 based on profile-based threading in combination with neighborhood effect. That is, distance- and orientation-based alignment further improves the alignment accuracy over the top template recognition performance achieved by profile-based threading with topological network neighborhood. The trend remains very similar considering the subset of 60 very hard CAMEO targets, although the exclusion of neighborhood effect results in a noticeable performance decline in this case (a mean TM-score drop from 0.335 to 0.329) that shows the effectiveness of incorporating topological network neighborhood for challenging threading scenarios. **Table 2** also reports a reference oracle method that uses TM-align (Zhang and Skolnick, 2005) to structurally align the experimental structure of the query protein with each of the templates in the template library to select the structurally closest template and the resulting optimal query-template alignment is then fed into the standard automodel() class of MODELLER to generate the 3D structures. Not surprisingly, the TM-align-based oracle achieves much better performance with a mean TM-score > 0.5 even for the very hard CAMEO targets (**Table 2**, last row), indicating that there is still a large room for improvement.

**Table 2.**
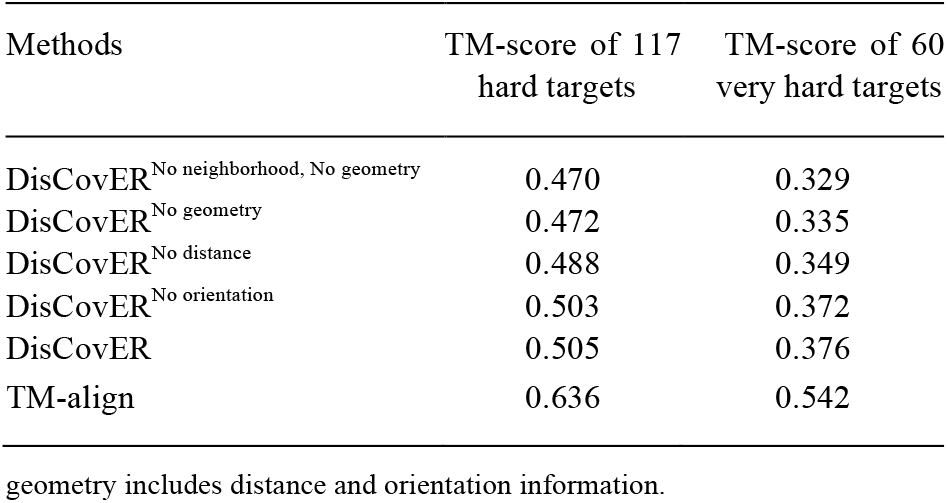
Contribution of individual features to DisCovER performance on CAMEO targets.

| Methods | TM-score of 117 hard targets | TM-score of 60 very hard targets |
| --- | --- | --- |
| DisCovER<sup>No neighborhood, No geometry</sup> | 0.470 | 0.329 |
| DisCovER<sup>No geometry</sup> | 0.472 | 0.335 |
| DisCovER<sup>No distance</sup> | 0.488 | 0.349 |
| DisCovER<sup>No orientation</sup> | 0.503 | 0.372 |
| DisCovER | 0.505 | 0.376 |
| TM-align | 0.636 | 0.542 |

geometry includes distance and orientation information.

### 3.3 3D model building using MODELLER from query-template alignment with additional restraints

To examine whether the additional information used in DisCovER for threading template selection and alignment can further improve the full-length 3D model building, we compare the standard automodel() class of MODELLER that builds 3D models using spatial restraints collected from query-template alignment to another approach using MODELLER that integrates additional restraints from predicted distances, orientations, and secondary structures. As shown in **Figure 2**, the mean TM-score attained by MODELLER with additional restraints is 0.544 over the 117 hard targets from CAMEO, better than that of standard MODELLER. MODELLER with additional restraints also attains 75 correct folds while shifting the TM-score distribution towards higher accuracy. That is, 3D model building using MODELLER from query-template alignment with additional restraints can be an effective use of the additional information used in DisCovER. We follow this model building approach henceforth.

**Fig 2.**
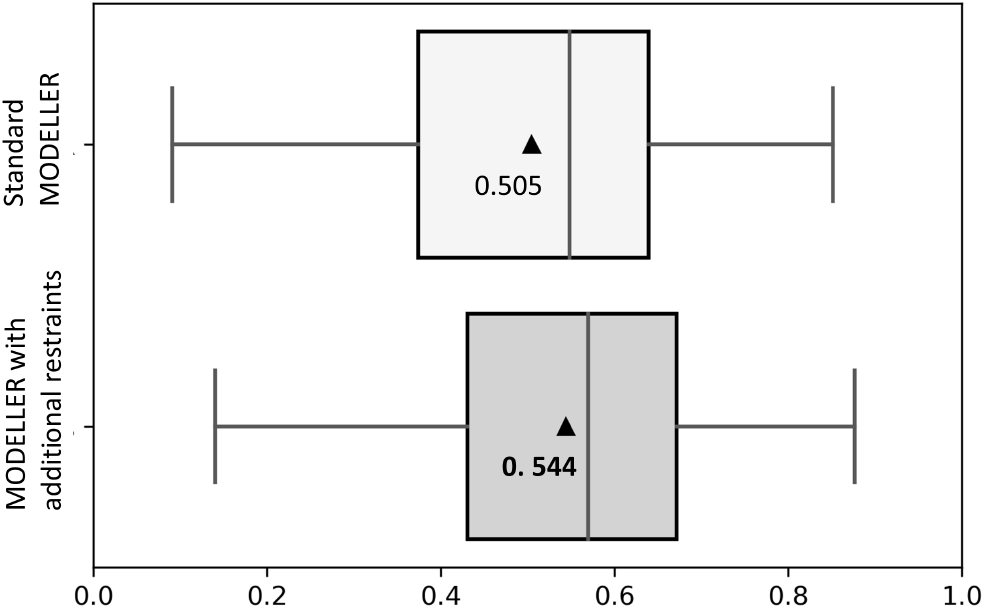
3D model building performance using standard MODELLER and MODELLER with additional restraints for 117 hard targets from CAMEO. The x-axis represents TM-score. The mean TM-scores are represented by triangles.

### 3.4 Performance comparison with CAMEO server RaptorX employing DeepThreader

We compare the performance of DisCovER to that of DeepThreader (Zhu *et al*., 2018) on the 60 very hard targets from CAMEO after downloading the predictions submitted by the CAMEO server RaptorX (Källberg *et al*., 2012; Zhu *et al*., 2018), which, according to the CAMEO assessment paper (Haas *et al*., 2019), employs the DeepThreader method, otherwise not publicly available to run. As shown in **Figure 3**, DisCovER achieves a mean TM-score of 0.435, outperforming RaptorX (mean TM-score of 0.397), while skewing the overall TM-score distribution towards higher accuracy. DisCovER attains higher TM-scores for 36 targets (60%) compared to RaptorX. In summary, DisCovER attains better overall performance over the 60 very hard targets from CAMEO for which HHpredB has TM-score less than 0.5. That is, DisCovER outperforms DeepThreader at one of the most challenging threading situations.

**Fig 3.**
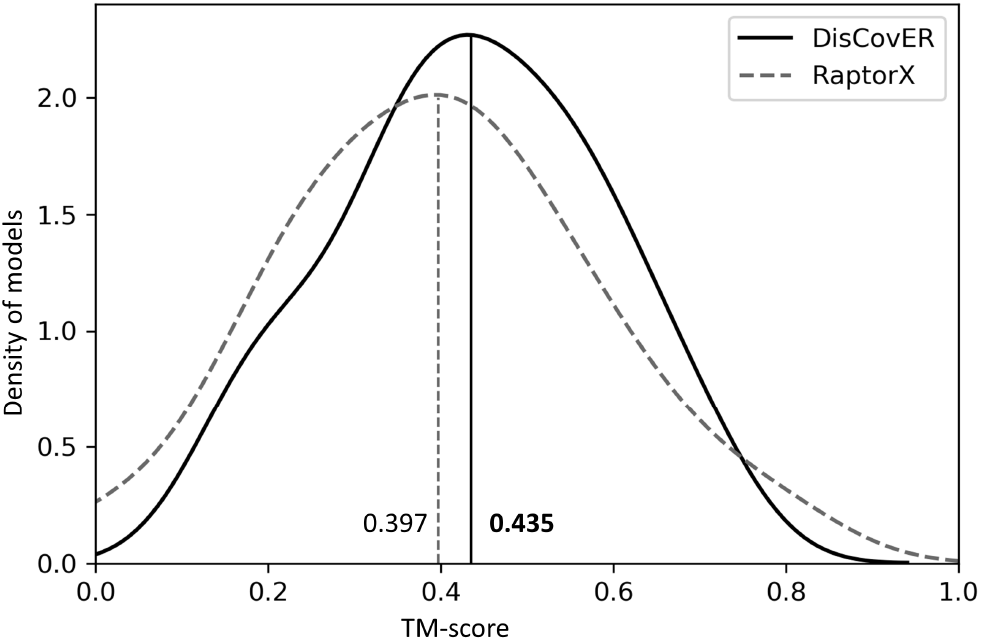
TM-score distribution of DisCovER and RaptorX on 60 very hard targets from CAMEO.

### 3.5 Performance on 131 hard targets from CATHER

The performance of DisCovER is further benchmarked against recent contact-assisted threading methods: CATHER (Du *et al*., 2020) and map_align (Ovchinnikov *et al*., 2017) by running DisCovER over 131 hard targets used in CATHER and directly comparing the mean TM-score with the results reported in the published work of CATHER (Du *et al*., 2020) over the same set. Consistent with our results on the CAMEO dataset, DisCovER significantly outperforms CATHER and map_align by attaining a mean TM-score of 0.551 as opposed to 0.456 of CATHER and 0.383 of map_align (**Supplementary Table S1**). DisCovER also greatly outperforms the reported mean TM-scores of other profile-based methods including HHpred (0.327), SparkX (0.349), and MUSTER (0.359) as well as the other contact-assisted approach EigenTHREADER (0.386). We note that we are unable to perform a target-by-target analysis since the CATHER paper reports only the average performance. Nonetheless, the better average performance of DisCovER continues to demonstrate its competitive advantage over current threading methods.

### 3.6 Effect of homologous information

The performance of DisCovER is weakly correlated with the number of effective sequence homologs, as quantified by Nf (Zhang *et al*., 2020). As shown in **Supplementary Figure S1**, the Spearman correlation between TM-score attained by DisCovER and Nf are 0.23 and 0.25 over the 117 hard targets from CAMEO and 131 hard targets from CATHER, respectively. In summary, there is a weak correlation between the performance of DisCovER and Nf.

### 3.7 Running time

**Figure 4** shows the running time of various threading methods with respect to the target length. All methods are run on the same Linux machine with 128 GB RAM and using a single CPU thread of Intel Xeon Processor (2.20 GHz). While it is expected that DisCovER is slower than most profile-based threading methods, but the running time of DisCovER is considerably faster than the top profile-based method CNFpred and orders of magnitude faster than the top contact-assisted approach CEthreader. Overall, DisCovER is reasonably efficient in terms of the running time.

**Fig 4.**
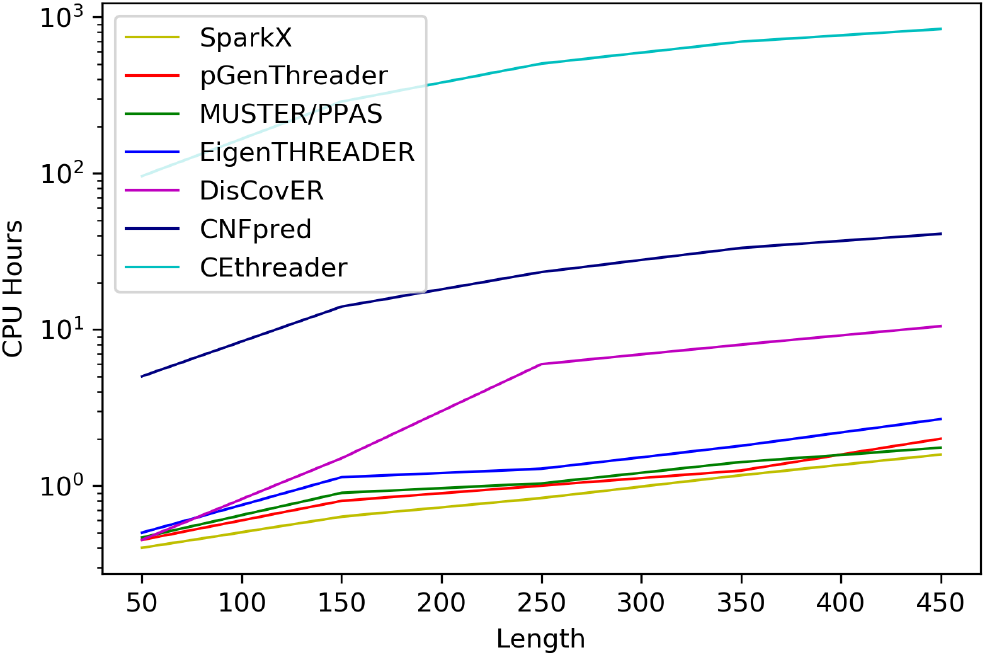
The running time of various threading methods with respect to the target length for 117 hard targets from CAMEO.

## 4 Conclusion

This article presents DisCovER, a new protein threading method that effectively integrates the covariational signal encoded in inter-residue distance and orientation information along with topological network neighborhood to significantly improve threading template selection and alignment for weakly homologous proteins. Experimental results show that our method yields better accuracy than existing threading methods, including profile-based methods and latest contact-assisted approaches such as CEthreader, EigenTHREADER, map_align, and CATHER. Controlled experiments reveal that distance and orientation information contributes significantly to the superior performance of DisCovER, complemented by the neighborhood effect particularly for weakly homologous proteins. At one of the most challenging threading situations, DisCovER outperforms the CAMEO server RaptorX employing the distance-based threading method DeepThreader.

It is important to note that the performance of our method is weakly correlated with the number of sequence homologs available for the query protein. This suggests that our distance- and orientation-based coevolutionary threading method DisCovER is well-suited for remotely homologous targets. Being reasonably efficient in terms of its running time, our study opens the possibility of successfully extending threading for many more protein sequences that were previously not amenable to template-based modeling.

## Acknowledgements

This work was made possible in part by a grant of high performance computing resources and technical support from the Alabama Supercomputer Authority and Auburn University Early Career Development Grant to DB.

## References

Altschul, S.F. et al. (1997) Gapped BLAST and PSI-BLAST: a new generation of protein database search programs. Nucleic Acids Res, 25, 3389–3402.

Berman, H.M. et al. (2000) The Protein Data Bank. Nucleic Acids Res, 28, 235–242.

Bhattacharya, S. and Bhattacharya, D. (2019) Does inclusion of residue-residue contact information boost protein threading? Proteins: Structure, Function, and Bioinformatics, 87, 596–606.

Buchan, D.W.A. and Jones, D.T. (2017) EigenTHREADER: analogous protein fold recognition by efficient contact map threading. Bioinformatics, 33, 2684– 2690.

Chen, C.-C. et al. (2019) TOPAS: network-based structural alignment of RNA sequences. Bioinformatics, 35, 2941–2948.

Dill, K.A. and MacCallum, J.L. (2012) The Protein-Folding Problem, 50 Years On. Science, 338, 1042–1046.

Du, Z. et al. (2020) CATHER: a novel threading algorithm with predicted contacts. Bioinformatics, 36, 2119–2125.

Greener, J.G. et al. (2019) Deep learning extends de novo protein modelling coverage of genomes using iteratively predicted structural constraints. Nat Commun, 10, 1–13.

Haas, J. et al. (2019) Introducing “best single template” models as reference baseline for the Continuous Automated Model Evaluation (CAMEO). Proteins: Structure, Function, and Bioinformatics, 87, 1378–1387.

Heffernan, R. et al. (2017) Capturing non-local interactions by long short-term memory bidirectional recurrent neural networks for improving prediction of protein secondary structure, backbone angles, contact numbers and solvent accessibility. Bioinformatics, 33, 2842–2849.

Henikoff, S. and Henikoff, J.G. (1994) Position-based sequence weights. Journal of Molecular Biology, 243, 574–578.

Jones, D.T. et al. (2015) MetaPSICOV: combining coevolution methods for accurate prediction of contacts and long range hydrogen bonding in proteins. Bioinformatics, 31, 999–1006.

Jones, D.T. et al. (2012) PSICOV: precise structural contact prediction using sparse inverse covariance estimation on large multiple sequence alignments. Bioinformatics, 28, 184–190.

Kabsch, W. and Sander, C. (1983) Dictionary of protein secondary structure: Pattern recognition of hydrogen-bonded and geometrical features. Biopolymers, 22, 2577–2637.

Källberg, M. et al. (2012) Template-based protein structure modeling using the RaptorX web server. Nature Protocols, 7, 1511–1522.

Kamisetty, H. et al. (2013) Assessing the utility of coevolution-based residue– residue contact predictions in a sequence- and structure-rich era. PNAS, 110, 15674–15679.

Li, Y. et al. (2019) ResPRE: high-accuracy protein contact prediction by coupling precision matrix with deep residual neural networks. Bioinformatics, 35, 4647–4655.

Lobley, A. et al. (2009) pGenTHREADER and pDomTHREADER: new methods for improved protein fold recognition and superfamily discrimination. Bioinformatics, 25, 1761–1767.

Ma, J. et al. (2012) A conditional neural fields model for protein threading. Bioinformatics, 28, i59–i66.

Ma, J. et al. (2013) Protein threading using context-specific alignment potential. Bioinformatics, 29, i257–i265.

Needleman, S.B. and Wunsch, C.D. (1970) A general method applicable to the search for similarities in the amino acid sequence of two proteins. Journal of Molecular Biology, 48, 443–453.

Ovchinnikov, S. et al. (2017) Protein structure determination using metagenome sequence data. Science, 355, 294–298.

Singh, R. et al. (2008) Global alignment of multiple protein interaction networks with application to functional orthology detection. PNAS, 105, 12763–12768.

Smith, T.F. and Waterman, M.S. (1981) Identification of common molecular subsequences. Journal of Molecular Biology, 147, 195–197.

Söding, J. (2005) Protein homology detection by HMM–HMM comparison. Bioinformatics, 21, 951–960.

Taylor, W.R. (1999) Protein structure comparison using iterated double dynamic programming. Protein Science, 8, 654–665.

Wang, S. et al. (2017) Accurate De Novo Prediction of Protein Contact Map by Ultra-Deep Learning Model. PLOS Computational Biology, 13, e1005324.

Webb, B. and Sali, A. (2014) Protein Structure Modeling with MODELLER. In, Kihara, D. (ed), Protein Structure Prediction, Methods in Molecular Biology. Springer, New York, NY, pp. 1–15.

Wu, Q. et al. (2020) Protein contact prediction using metagenome sequence data and residual neural networks. Bioinformatics, 36, 41–48.

Wu, S. and Zhang, Y. (2007) LOMETS: A local meta-threading-server for protein structure prediction. Nucleic Acids Res, 35, 3375–3382.

Wu, S. and Zhang, Y. (2008) MUSTER: Improving protein sequence profile–profile alignments by using multiple sources of structure information. Proteins: Structure, Function, and Bioinformatics, 72, 547–556.

Xu, J. (2019) Distance-based protein folding powered by deep learning. PNAS, 116, 16856–16865.

Xu, J. and Wang, S. (2019) Analysis of distance-based protein structure prediction by deep learning in CASP13. Proteins: Structure, Function, and Bioinformatics, 87, 1069–1081.

Xu, J. and Zhang, Y. (2010) How significant is a protein structure similarity with TM-score = 0.5? Bioinformatics, 26, 889–895.

Yang, J. et al. (2020) Improved protein structure prediction using predicted interresidue orientations. PNAS, 117, 1496–1503.

Yang, J. et al. (2015) The I-TASSER Suite: protein structure and function prediction. Nature Methods, 12, 7–8.

Yang, Y. et al. (2011) Improving protein fold recognition and template-based modeling by employing probabilistic-based matching between predicted one-dimensional structural properties of query and corresponding native properties of templates. Bioinformatics, 27, 2076–2082.

Zhang, C. et al. (2020) DeepMSA: constructing deep multiple sequence alignment to improve contact prediction and fold-recognition for distant-homology proteins. Bioinformatics, 36, 2105–2112.

Zhang, Y. and Skolnick, J. (2004) Scoring function for automated assessment of protein structure template quality. Proteins: Structure, Function, and Bioinformatics, 57, 702–710.

Zhang, Y. and Skolnick, J. (2005) TM-align: a protein structure alignment algorithm based on the TM-score. Nucleic Acids Res, 33, 2302–2309.

Zheng, W. et al. (2019) Detecting distant-homology protein structures by aligning deep neural-network based contact maps. PLOS Computational Biology, 15, e1007411.

Zhu, J. et al. (2018) Protein threading using residue co-variation and deep learning. Bioinformatics, 34, i263–i273.

