## Supplementary Information for "DisCovER: distance- and orientation-based covariational threading for weakly homologous proteins"

### Supplementary Information
*for*

### DisCovER: distance- and orientation-based covariational threading for weakly homologous proteins

Sutanu Bhattacharya[1], Rahmatullah Roche[1], Debswapna Bhattacharya [1,2,*]

[1]Department of Computer Science and Software Engineering, [2]Department of Biological Sciences, Auburn University, AL 36849, USA.

*To whom correspondence should be addressed.

**Supplementary Table. S1.** Benchmark results on 131 hard targets from CATHER. Average TM-score of first-ranked 3D full-length models are evaluated.

| HHpred | SparkX | MUSTER | map_align | EigenTHREADER | CATHER | DisCovER |
|---|---|---|---|---|---|---|
| 0.327 | 0.349 | 0.359 | 0.383 | 0.386 | 0.456 | **0.551** |

*Note:* Except DisCovER, the results of other methods are taken from the reported results of CATHER for the targets classified as "hard" by CATHER. Templates with sequence identity >30% to the query protein are excluded. The value in bold represents the best performance.

**Supplementary Fig. S1.** TM-score of DisCovER *vs* Nf (number of sequence homologs) on (A) 117 hard targets from CAMEO having a Spearman correlation of 0.23, (B) 131 hard targets from CATHER having a Spearman correlation of 0.25.

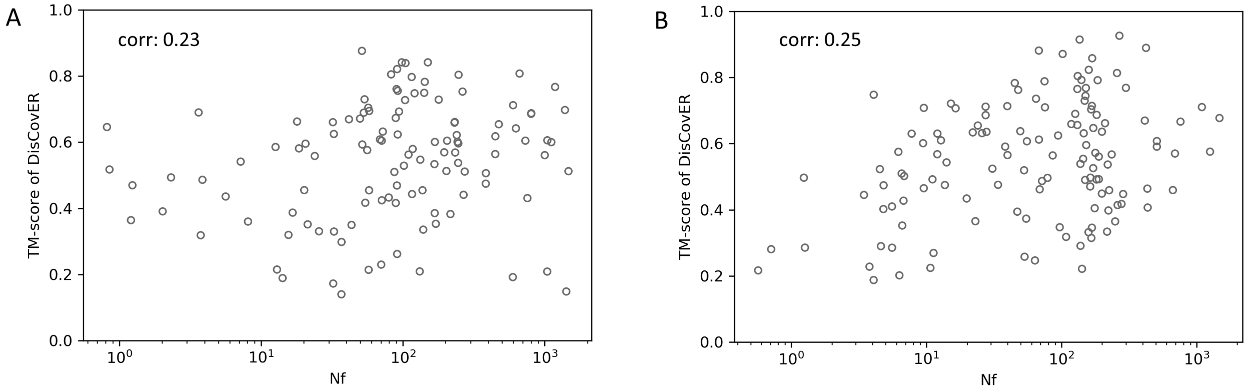
